# Data-independent acquisition coupled with electron-activated dissociation for in-depth structure elucidation of fatty acid ester of hydroxy fatty acids

**DOI:** 10.1101/2024.12.11.627939

**Authors:** Yuto Kurizaki, Yuki Matsuzawa, Mikiko Takahashi, Hiroaki Takeda, Mayu Hasegawa, Makoto Arita, Junki Miyamoto, Hiroshi Tsugawa

## Abstract

Fatty acid esters of hydroxy fatty acid (FAHFAs) are a biologically important class of lipids known for their anti-inflammatory and anti-diabetic effects in animals. The physiological activity of FAHFAs varies depending on the length of the carbon chain, number and position of double bonds (DBs), and the position of the hydroxyl (OH) group. Moreover, gut bacteria produce FAHFAs with more diverse structures than those produced by the host, which necessitates a FAHFA-lipidomics approach grasping their diverse structures to fully understand the physiological and metabolic significance of FAHFAs. In this study, we developed a methodology for in-depth structural elucidation of FAHFAs. First, FAHFAs were enriched using a solid-phase extraction (SPE) system coated with titanium and zirconium dioxide, which separated these analytes from neutral lipids and phospholipids. The fractionated metabolites were then derivatized using *N*,*N*-dimethylethylenediamine (DMED) to facilitate FAHFA detection in the positive ion mode of a liquid chromatography-tandem mass spectrometry (LC-MS/MS) system. A data-independent acquisition technique known as sequential window acquisition of all theoretical mass spectra (SWATH-DIA) was used to collect sequential MS/MS spectra of the DMED-derivatized fatty acid metabolites. Structural elucidation was based on the fragment ions generated by electron-activated dissociation (EAD). DMED-FAHFAs were annotated using the newly updated MS-DIAL program, and FAHFA isomers were quantified using the MRMPROBS program, which quantifies lipids based on SWATH-MS/MS chromatograms. This procedure was applied to profile the FAHFAs present in mouse fecal samples, characterizing seven structures at the molecular species level, 63 structures at the OH position-resolved level, and 15 structures at both the DB and OH position-resolved levels using the MS-DIAL program. In the MRMPROBS analysis, 2OH and 3OH hydroxy fatty acids with more than 20 carbon atoms were predominantly expressed, while 5OH–13OH hydroxy fatty acids with 16 or 18 carbon atoms were the major components, abundant at positions 5, 7, 9, and 10. Furthermore, age-related changes in FAHFA isomers were also observed, where FAHFA 4:0/2O(FA 26:0) and FAHFA 16:0/10O(FA 16:0) significantly increased with age. In conclusion, our study offers a novel LC-SWATH-EAD-MS/MS technique with the updates of computational MS to facilitate in-depth structural lipidomics of FAHFAs.

**TOC graphics:** 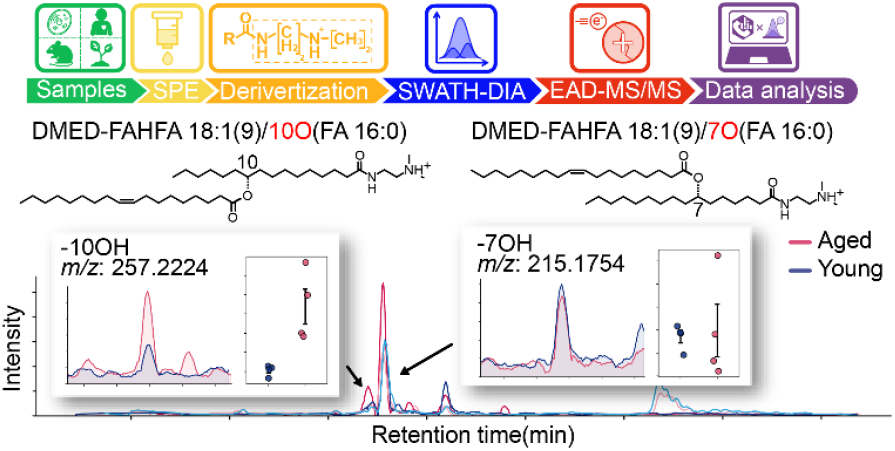

---

Fatty acid esters of hydroxy fatty acid (FAHFAs) are defined as fatty acyls produced by various organisms, including microbes, plants, and animals^1^. In humans, FAHFAs are important endogenous metabolites with anti-diabetic and anti-inflammatory effects^2^. These molecules are biosynthesized in humans and mice by esterifying the carboxylic acid of a fatty acid with the hydroxyl group of a hydroxy fatty acid, in the presence of PNPLA2 (patatin-like phospholipase domain containing 2), also known as adipose triglyceride lipase (ATGL)^3^. The structure of FAHFAs is defined by the number of carbons in the fatty acid chain, number and position of double bonds, and position of the hydroxyl group. Importantly, their physiological activities often vary with their structures. For example, palmitic acid esters of hydroxystearic acid (PAHSA) represent a prominent FAHFA family found in the mammalian blood and white adipose tissue (WAT)^2^. Among the PAHSAs, the blood levels of 5-PAHSA significantly decreased in patients with insulin resistance, whereas 9-PAHSA substantially improved glucose tolerance and had anti-inflammatory effects. A study investigating insulin secretion, glucose uptake, and anti-inflammatory effects showed that glucose-stimulated insulin secretion was enhanced by branched FAHFA isomers containing unsaturated fatty acids with ester linkages at a higher carbon position^4^. In contrast, FAHFAs branched at lower carbon positions from the carboxylate head group exhibited more consistent anti-inflammatory effects than those branched at higher carbon positions. Additionally, gut bacteria produce unique FAHFAs, particularly those involving ester bonds between short-chain fatty acids and 2-hydroxy fatty acids^5^. These FAHFAs are referred to as short-chain FAHFAs (SCFAHFAs), and an acyl alpha-hydroxy fatty acid (AAHFA) lipid class^6,7^. While short-chain fatty acids produced by gut bacteria are possibly important for the homeostasis of intestinal immunity, whether these SCFAHFAs function in short-chain fatty acid storage or the SCFAHFA structure itself plays a significant role is not yet clear. Since FAHFA originate from not only the host, but also gut bacteria and dietary nutrients, humans interact with various FAHFAs.

Liquid chromatography coupled with tandem mass spectrometry (LC-MS/MS) is widely used for FAHFA metabolic profiling. While several amino acid conjugates of FAHFAs, known as *N*-acyl amides, can be characterized in the positive ion mode of electron spray ionization (ESI), normal FAHFA molecules with carboxylic acid moieties are detected in the negative ion mode^8^. In addition, multistage fragmentation using the MS3 technique revealed the hydroxyl position of the FAHFAs^9^. Derivatization methods have also been used in FAHFA profiling to increase the sensitivity of metabolite screening^10-12^. A well-known method involves the use of *N*,*N*-dimethylethylenediamine (DMED), which converts the carboxylic acid moiety to a tertiary amine, thus detecting DMED-FAHFA in positive ion mode^11,12^. The use of deuterated DMED (such as d4-DMED) facilitates relative quantification between sample groups when an appropriate control group is established^12^. In addition, DMED derivatization is applicable to not only FAHFAs, but also any fatty acyls containing carboxyl groups, where free fatty acids and hydroxy fatty acids can also be characterized.

In this study, we developed a novel analytical technique to accelerate the in-depth structural elucidation of FAHFAs by incorporating electron-activated dissociation (EAD), data-independent acquisition (DIA), and sequential window acquisition of all theoretical fragment ion spectra (SWATH-DIA). Although collision-induced dissociation (CID) use accelerates the characterization of fatty acid and hydroxy fatty acid pairs in FAHFA^11^, the CID-MS/MS spectrum does not provide information on the hydroxy and double bond positions unless additional techniques such as MS3 are incorporated. In contrast, the ESI(+)-EAD-MS/MS spectrum of DMED-FAHFA generated ions specific to the hydroxy position. Furthermore, the positions of the double bonds in highly abundant FAHFAs may be estimated. To maximize the advantages of this analytical chemistry technique, we optimized the sample preparation and data analysis. Because FAHFAs and other fatty acid metabolites are present in smaller amounts than phospholipids and neutral lipids such as triacylglycerols, fatty acid metabolites from biological samples should be fractionated and enriched. In a recent study, we found that using a solidphase extraction (SPE) column with titanium and zirconium dioxide under optimal solvent conditions fractionated fatty acids from phospholipids and neutral lipids^13^. Thus, we applied SPE to fractionate and enrich FAHFAs. In addition, we developed an algorithm to annotate the EAD-MS/MS spectra of DMED-FAHFA by improving MS-DIAL. Since the EAD-MS/MS of DMED-FAHFA shows high sensitivity for detecting fragments dependent on the hydroxy position, we proposed to quantify FAHFA isomers at the hydroxy position using the SWATH-MS2 chromatogram with the MRMPROBS software program^14,15^. Essentially, we established a data analysis pipeline for OH-position-resolved FAHFA profiling by screening molecular species in MS-DIAL based on the acquired SWATH-DIA data and then quantifying the MS2 chromatogram of the hydroxy position isomers in MRMPROBS. The optimized procedure was applied to fecal lipidomics in young and aged mice.

## EXPERIMENTAL SECTION

### Chemical regents

Sixteen FAHFA standards were purchased from Cayman Chemicals (Ann Arbor, Michigan, USA) (**Supplementary Table 1**). Ammonium acetate solution and chloroform (CHCl_3_) for high-performance LC (HPLC), ethylenediaminetetraacetic acid (EDTA), acetonitrile (ACN), methanol (MeOH), 2-propanol (IPA), and ultrapure water (H_2_O) of quadrupole time-of-flight MS (QTOF-MS) grade were purchased from FUJIFILM Wako Pure Chemical Corp. (Osaka, Japan). Methyl tert-butyl ether (MTBE) for HPLC was purchased from Sigma-Aldrich (Tokyo, Japan). Triethylamine (TEA), 2-chloro-1-methylpyridinium iodide (CMPI), and *N*,*N*-dimethylethylenediamine (DMED) for lipid derivatization were purchased from Hayashi Pure Chemical Ind. (Osaka, Japan), Combi Blocks Inc. (CA, USA), and Tokyo Chemical Industry Co. (Tokyo, Japan), respectively. MonoSpin Phospholipid for solid phase extraction (SPE) was obtained from GL Sciences Inc. (Tokyo, Japan) for the monolithic silica column coated TiO_2_ and ZrO_2_ (φ 4.2 μm × 1.5 mm).

### Animal experiment

The animal experiments were conducted in accordance with the ethical protocol approved by the Tokyo University of Agriculture and Technology (R5-50). C57BL/6J male mice were purchased from SLC (Shizuoka, Japan). The mice were fed CE-2 chow (CLEA Japan, Tokyo, Japan). Feces from young (11-week-old) and aged (24-month-old) mice were harvested, immediately frozen, and stored at −80 °C until lipid extraction.

### Standard mixture analysis

A mixture of 16 FAHFA standards dissolved in MeOH was prepared, and the concentration of each metabolite was adjusted to 1 μM. The initial mixture was diluted by factors of 2, 5, 10, 20, 50, and 100 with MeOH.

### Lipid extraction

Fecal samples were homogenized using a multi-bead shocker with a metal cone (YASUI KIKAI, Japan) at 2500 rpm for 15 s and 1 mL MeOH was added. MeOH containing 10 mg fecal sample was transferred to a new 2.0 mL tube, and additional MeOH was added to bring the total volume to 225 μL. After adding 750 μL MTBE, 188 μL H_2_O was added, shaken for 10 min, and centrifuged at 16,000 rpm for 3 min. Then, 600 μL upper organic phase was collected in a clean tube and dried using a centrifuge evaporator. For analysis without SPE technique, the dried sample was resuspended in 30 μL MeOH containing 1 μM 9-PAHSA (d9) internal standard, and transferred into LC-MS vials (Agilent Technologies, Santa Clara, CA, USA). Lipid fractionation was performed as described in a previous report^13^. The dried lipid extract was dissolved in 100 μL CHCl_3_. A MonoSpin Phospholipid SPE column was activated with 200 μL CHCl_3_ before loading the lipid extract. The lipid extract was then applied to the SPE column. After centrifuging at 300× *g* and 4 °C for 1 min to elute neutral lipids, 200 μL MeOH/HCOOH (99:1, v/v) was applied to the SPE columns and then centrifuged at 300× *g* and 4 °C for 3 min to elute the fatty acid metabolites including FAHFAs. The fatty acid fraction was dried and resuspended in 30 μL MeOH containing 1 μM 9-PAHSA (d9) internal standard. For analysis of the DMED-derivatized metabolites, the dried samples were derivatized without resuspension.

### FAHFA derivatization

The dried sample was resuspended with the mixture of 200 μL ACN, 15 μL CMPI, and 30 μL TEA. After voltex for 20 s, the solvent was incubated for 5 min at 40 °C using block incubator BI-535 (ASTEC, Japan). After 30 μL DMED was added to the solvent, the sample was incubated for 1 h at 40 °C. The solvent was dried up and resuspeneded in 30 μL MeOH containing 1 μM 9-PAHSA (d9) internal standard.

### LC-MS measurement

The LC system was an ExonLC system (SCIEX, Framingham, MA, USA). Lipids were separated on an ACQUITY UPLC CSH C18 Column (100×2.1 mm^2^: 1.7 µm) (Waters, U.S.A). The column was maintained at 40 °C and 0.2 mL/min flow rate. The mobile phase comprised (A) 9:9:2 (v/v/v) ACN:MeOH:H_2_O with ammonium acetate (5 mM) and 0.1% acetic acid, and (B) 9:1 (v/v) IPA:ACN with 5 mM ammonium acetate and 0.1% acetic acid. A sample volume of 3 μL was injected. The separation was performed under the following gradient: 0 min 10% (B), 5.0 min 45% (B), 15.0 min 55% (B), 20.0 min 90% (B), 21 min 90% (B), 21.01 min 10% (B), 26.0 min 10% (B). The temperature of the rack was maintained at 4 °C. The MS detection of lipids was performed using QTOF-MS (ZenoTOF 7600; SCIEX, Framingham, MA, USA). The parameters for ESI(+)- and ESI(−)-DDA-CID analyses were as follows: MS1 and MS2 mass ranges, *m/z* 75–1250; MS1 accumulation time, 200 ms; Q1 resolution, units; MS2 accumulation time, 50 ms; maximum candidate ions, 10; ion source temperature, 450 °C; and CAD gas, 7. The following settings were used for positive/negative ion mode, independently: ion source gas 1, 40/50 psi; ion source gas 2, 80/50 psi; curtain gas, 30/35 psi; spray voltage, 5500/−4500 V; declustering potential, 80/−80 V; and collision energy, 40/−42 ± 15 eV. The parameters for ESI(+)-DDA-EAD were as follows: MS2 mass range, *m/z* 140– 1250; MS2 accumulation time, 100 ms; maximum number of candidate ions, eight; kinetic energy, 14 eV; and EAD RF, 140. The other settings were the same as those used for CID. The parameters for the ESI(+)-SWATH-DIA-EAD analysis were as follows: MS1 accumulation time, 100 ms; MS2 accumulation time, 50 ms; Q1 window, 25 Da; and precursor scanning range, *m/z* 300–700. The other settings were the same as those used for ESI(+)-DDA-EAD. In addition, the EAD-MS/MS spectral patterns of authentic FAHFA standards were confirmed by variable kinetic energies involving one CID setting (45 eV) and eight kinetic energies (8, 10, 12, 14, 16, 18, and 20 eV KE) acquired in the targeted product ion scanning mode, known as MRM-HR in SCIEX. An isocratic mode of 80% B was used by the same solvent properties with an InertSustainSwift C18 column (30×2.1 mm^2^, 3 μm; GL Sciences, Japan) maintained at 40 °C with 0.2 mL/min flow rate.

## RESULTS AND DISCUSSION

### Strategy of FAHFA profiling using LC-DIA-EAD-MS/MS

Here, we propose an in-depth and comprehensive structural elucidation strategy for FAHFAs (**Figure 1**). We investigated the EAD-MS/MS patterns of DMED-FAHFA using authentic standards and developed an environment for the automatic annotation of DMED-FAHFA molecules using MS-DIAL 5. For biological sample analysis, we applied an SPE-based fractionation method to enrich fatty acid metabolite molecules separated from glycerolipids and glycerophospholipids. The derivatized fatty acid metabolites were analyzed using LC-SWATH-DIA-EAD-MS/MS and annotated using the MS-DIAL 5 program. As MS-DIAL uses an MS1-based extracted ion chromatogram (EIC) to quantify the biomolecules, the co-eluted structural isomers were not distinguished. Thus, the potential structural candidates in the chromatographic peaks were further considered in the MRMPROBS program, where the product ion trace of a unique fragment ion specific to the OH position was used for quantification.

**Figure 1.**
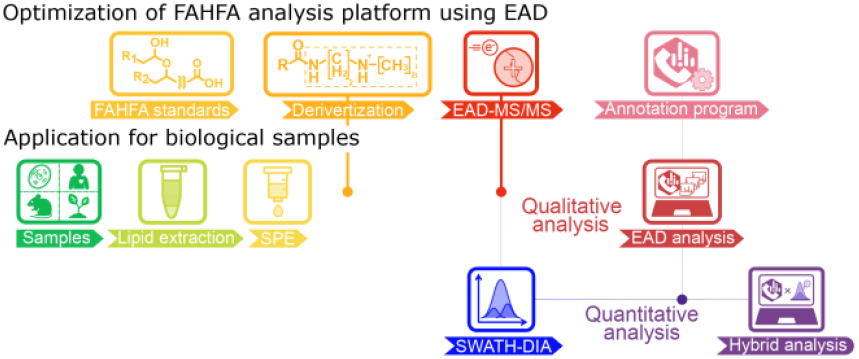
A strategy for in-depth profiling of fatty acid esters of hydroxy fatty acid (FAHFAs) with electron-activated dissociation (EAD). Elucidation of the derivatized FAHFA MS/MS spectral patterns were elucidated using authentic standards. The optimized workflow was applied to biological samples, where a solid-phase extraction (SPE) method is employed to enrich FAHFA molecules. The EAD-associated sequential window acquisition of all theoretical mass spectra (EAD-SWATH-DIA) data were processed by MS-DIAL for characterizing FAHFA metabolites, and MRMPROBS was used for SWATH-DIA-MS2 chromatogram-based lipid quantification.

### Evaluating the EAD-MS/MS spectrum pattern of FAHFAs

We analyzed the DMED-derivatized forms of 16 authentic standard FAHFAs using ESI(+)-CID-MS/MS and ESI(+)-EAD-MS/MS; the fragmentation pattern was exemplified by 12-oleic acid ester of HSA (12-OAHSA; FAHFA 18:1(9Z)/18:0;12OH) (*m/z* 635.6085; C_40_H_79_N_2_O_3_^+^) (**Figure 2a**). Fragments of H_2_O loss from DMED-HFA (*m/z* 353.3526; C_22_H_45_N_2_O^+^) and H_2_O and C_2_H_7_N losses from DMED-HFA (*m/z* 308.2948; C_20_H_38_NO^+^) were detected as the major fragment ions in both CID- and EAD-MS/MS. Two fragments with a 45 Da difference can be used to detect the presence of the DMED moiety in complex lipids. Furthermore, the fragment ion, which considers the cleavage of the branched chain esterified bond and carbon–carbon bond at the branched position (*m/z* 285.2537; C_16_H_33_N_2_O_2_), was also detected as an abundant ion in EAD-MS/MS, which is a unique fragment in EAD and absent in CID, indicating the OH position of HFA. In addition, the ion abundance increases of the hydrogen loss fragment ion (H-loss) and the radical fragment ion (radical) by the cleavages of C11– C12 and C7–C8 bonds, respectively, in the FA chain were observed in the EAD-MS/MS. The V-shaped pattern, whose valley corresponds to the C=C position, is the diagnostic criterion for interpreting the C=C position. The increased abundance of H-loss and radical ions is due to fragment ion stabilization by McLafferty rearrangement and allyl radical formation, respectively^16^. Unique fragmentation patterns for the determination of the OH- and C=C positions in DMED-FAHFA were observed for all authentic standards examined in this study. Furthermore, we confirmed that 14 eV kinetic energy provided higher sensitivities for product ions specific to the –OH and C=C positions than those of the other conditions (**Figure 2b**). However, the fragment ions reflecting the double-bond position had <5% relative intensity ratio and the limit of detection (LOD) for doublebond-specific ions was 200 nM, even under ideal conditions using standards (**Figure 2c**). While V-shaped pattern recognition may be effective when only branched chain fatty acids, such as OAHSA, have double bonds, it becomes difficult to interpret spectra when polyunsaturated fatty acids are involved or when both FA and HFA contain double bonds. Therefore, this study focused primarily on profiling the OH-position isomers of FAHFA, and C=C positional information was assigned only when we were confident in interpreting the double-bond position-specific fragment ions.

**Figure 2.**
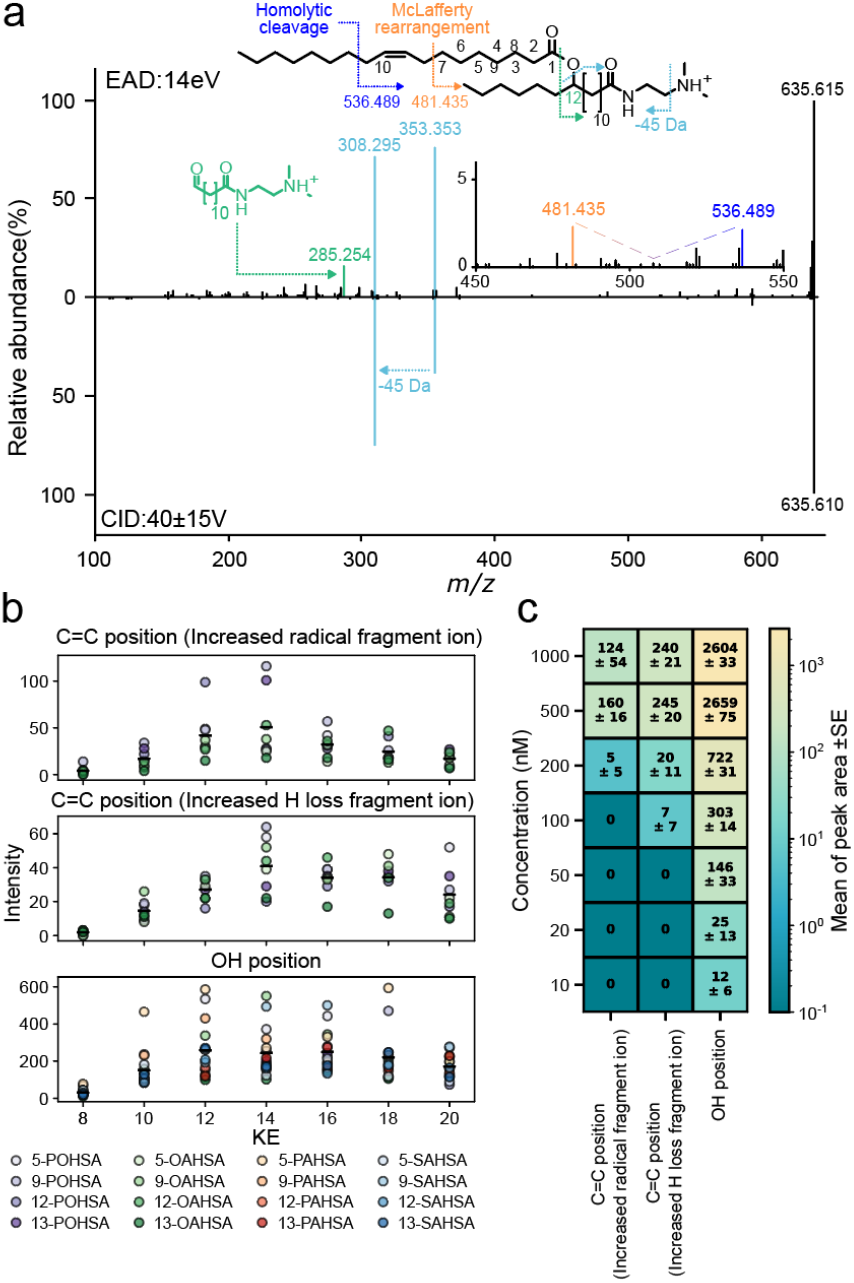
Elucidation of *N*,*N*-dimethylethylenediamine-derivatized fatty acid ester of hydroxy fatty acid (DMED-FAHFA) fragment ions for lipid annotation and quantification. (a) The MS/MS spectra of DMED-derivatized 12-oleic acid ester of hydroxy stearic acid (12-OAHSA) standard using electron-activated dissociation (EAD; 14 eV, upper panel) and collision-induced dissociation (CID; 40±15V, lower panel). The V-shaped product ion pattern was also described by zooming at the range from *m/z* 450 to 550. (b) Product ion abundances of diagnostic fragment ions including two C=C position-related ions (radical and H-loss) and OH position-related ion for various FAHFA isomers. The abundance of C=C position-resolved ion is increased by fragment ion stabilization by the McLafferty rearrangement for the hydrogen loss (H-loss) fragment ion or allyl radical formation for the radical fragment ion (radical). (c) Heatmap showing the limits of detection for the diagnostic ions from various FAHFA molecular species at 10–1000 nM. Values represent the mean MS/MS chromatogram peak area ± standard error (SE) (n=3).

### Evaluation of the SPE method to enrich FAHFA molecules

We have recently developed a simple method for fractionating lipid components using SPE columns based on titanium and zirconium dioxide^13^. This method separates lipids into (1) neutral lipids, such as sterol esters and triacylglycerols; (2) phospholipids; and (3) other lipids, whose representative subclasses are fatty acids, ceramides, and glycolipids. By removing the abundant neutral lipids and phospholipids, fatty acids, including FAHFA, can be enriched, which improves the sensitivity of FAHFAs profiling. We applied the SPE procedure to mouse fecal samples, resulting in the fractionation of free fatty acids and FAHFAs with a high recovery rate of 100% in the fraction eluted with methanol (MeOH) containing 1% formic acid (**Figure 3** and **Supplementary Figure 1**).

**Figure 3.**
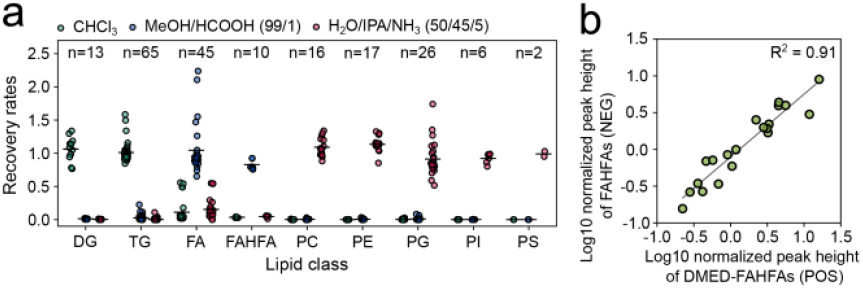
Lipid recovery rates using solid-phase extraction and abundance correlation of fatty acid esters of hydroxy fatty acid (FAHFAs) with or without *N*,*N*-dimethylethylenediamine (DMED) derivatization. (a) Recovery rates of various lipid classes (DG, TG, FA, FAHFA, PC, PE, PG, PI, PS) in three fractions: CHCl_3_, MeOH/HCOOH (99/1), and H_2_O/IPA/NH_3_ (50/45/5) with titanium and zirconium dioxide-coated solid phase extraction (SPE) method. The recovery rate was calculated by the peak heights between non-SPE and SPE. Each dot represents the recovery rate of a molecule in the lipid subclass. The number in the parentheses denotes the number of detected lipids in mouse fecal samples. (b) Scatter plot showing the correlation between log10 normalized peak heights of non-derivatized FAHFAs detected in negative ion mode (NEG) and DMED-derivatized FAHFAs detected in positive ion mode (POS). The R-square value (R^2^ = 0.91) was also described.

In the experiment in which we concentrated the fractionated FAHFA metabolites 5, 10, and 30 times, the number of characterized FAHFAs with essential fragment ions for fatty acid composition and OH-position determination increased from 20 (no enrichment; control) to 32 (5 times), 49 (10 times), and 77 (30 times), respectively (**Supplementary Figure 2**). Furthermore, a correlation plot of quantitative values from the positive ion mode results of DMED-FAHFA after fractionation and derivatization and from the negative ion mode results of FAHFA without fractionation and derivatization showed high R-squre value (R^2^=0.91), where the FAHFA metabolites detected in the negative ion mode were evaluated (**Figure 3b**). However, the ion abundances of DMED-FAHFAs are not compatible with those of the non-derivatized forms analyzed in the negative ion mode. Normalization using a proper internal standard is needed, and deuterium-labeled FAHFA (9-PAHSA-d9) was used in this study.

### Using SWATH-DIA for profiling co-eluted FAHFAs

We confirmed the elution times of the OH-positional isomers of palmitoleic acid ester of hydroxy stearic acid (POHSA), PAHSA, OAHSA, and stearic acid ester of HSA (SAHSA) under the 30 min LC condition used in this study (**Figure 4a**). The results showed that the peaks of FAHFAs with significantly different hydroxyl positions, such as 5-OAHSA and 12-OAHSA, were separated, whereas the peaks of FAHFAs with only one OH position difference, such as 12-OAHSA and 13-OAHSA, had almost identical retention times. Therefore, we used the SWATH-DIA technique to characterize the OH-positional isomers of FAHFAs, in which the product ion traces (MS/MS chromatograms) of the OH-position-specific fragment ions were sequentially monitored and used for lipid quantification. The precursor window size for SWATH-DIA was fixed at 25 Da, and the scan range for precursor selection was set from 300 Da to 700 Da, covering DMED-FA 16:0 to DMED FAHFA 40:0. The accumulation time for MS/MS monitoring of the precursor window was fixed at 50 ms, with a total cycle time including a full MS scan, of 0.9 seconds. Under this cycle time, approximately 20 data points were obtained for each peak under our LC conditions.

**Figure 4.**
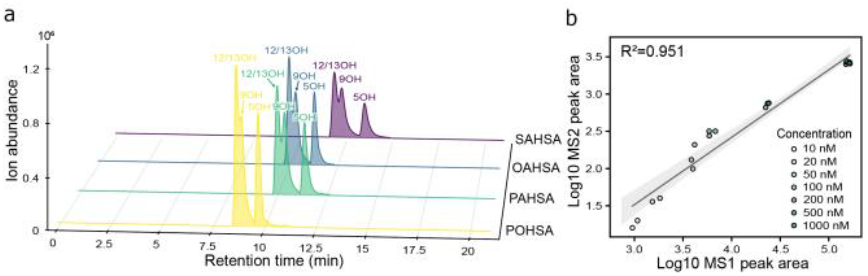
Chromatographic separation and quantitative analysis of fatty acid esters of hydroxy fatty acid (FAHFA) isomers. (a) Extracted ion chromatograms (EICs) of FAHFA isomers after DMED derivatization. The 3D chromatogram displays retention time (min) on the x-axis, intensity on the y-axis, and different FAHFA types (POHSA, PAHSA, OAHSA, SAHSA) on the z-axis. Peaks for various OH position isomers (5OH, 9OH, 12/13OH) are labeled for each FAHFA isomer. (b) Correlation between MS1 and MS2 peak areas for 12-OAHSA. The scatter plot shows the relationship between log10-transformed MS1 peak areas (x-axis) obtained from MS-DIAL analysis and log10-transformed MS2 peak areas (y-axis) exported by MRMPROBS, showing 0.951 R-square value.

MS1 and MS2 spectral data from the SWATH-DIA method were used to annotate and quantify the FAHFA metabolites. In our previous study, we showed that MS-DIAL maximizes the value of SWATH-DIA data for metabolite annotation, and MRMPROBS maximizes the value for metabolite quantification, where co-eluted isomers can be distinguished by unique product ions^17-20^. MS-DIAL provides putative FAHFA candidates for a single chromatographic peak that may have co-eluted molecules based on MS/MS spectral information, and the MRMPROBS reference library containing information on co-eluted lipid metabolites was automatically exported from the MS-DIAL program. This library file contains information on quantitative and qualitative product ions for targeted FAHFA metabolites, as well as intensity ratios at the top peak positions among the diagnostic ions, together with retention time information, allowing accurate qualitative analysis in MRMPROBS. We confirmed the relationship between the quantification values of the full MS1 precursor ion transitions from MS-DIAL and the MS2 product ion traces from MRMPROBS using authentic standards, which showed high correlation coefficients (R2=0.951) (**Figure 4b**). Although this method provides relative quantification values among samples, there is a possibility of over-or underestimation of lipid abundance among samples because the matrix effect varies among samples. Thus, careful interpretation of the results should be performed, and if necessary, quantitative values should be validated using another or-thogonal method.

### FAHFA metabolic alterations between young and aged mice

We applied our experimental procedure using computational mass spectrometry to fecal samples obtained from young and aged mice (**Figure 5a**). A total of 85 FAHFAs were characterized, and the OH positions and both the C=C and OH positions were resolved for 63 and 15 molecules, respectively (**Supplementary Table 2**). In addition, free HFA were also characterized in the same EAD-MS/MS dataset as the DMED-HFA form (**Supplementary Figure 3**). We further characterized the OH-positional isomers of 38 lipid metabolites, including HFA and FAHFA, using MRMPROBS: these 38 metabolites were selected based on the signal-to-noise ratios >10 and the fact that saturated fatty acids are present as HFA, where the confidence of the OH-specific fragment ion in annotation is increased when compared to FAHFAs containing unsaturated fatty acids as HFA. The results showed that FAHFAs containing long-chain saturated HFA (LHFA), which have more than 20 carbons, were abundant in feces compared to FAHFA 18:1/18:0;O, FAHFA 16:0/16:0;O, and FAHFA 18:2/16:0;O, which are biosynthesized by mammalian cells. Our results indicate that 7OH-FA 16:0 was the most abundant among FAHFA isomers containing HFA 16:0, while 10OH-FA 18:0 was abundant in HFA 18:0 containing FAHFA isomers. In contrast, FAHFAs with LHFA mostly contain SCFAs as branched chains. According to our previous and related results, SCFAHFAs are mostly absent in germ-free and antibacterial drug-treated mice^7,21^.

**Figure 5.**
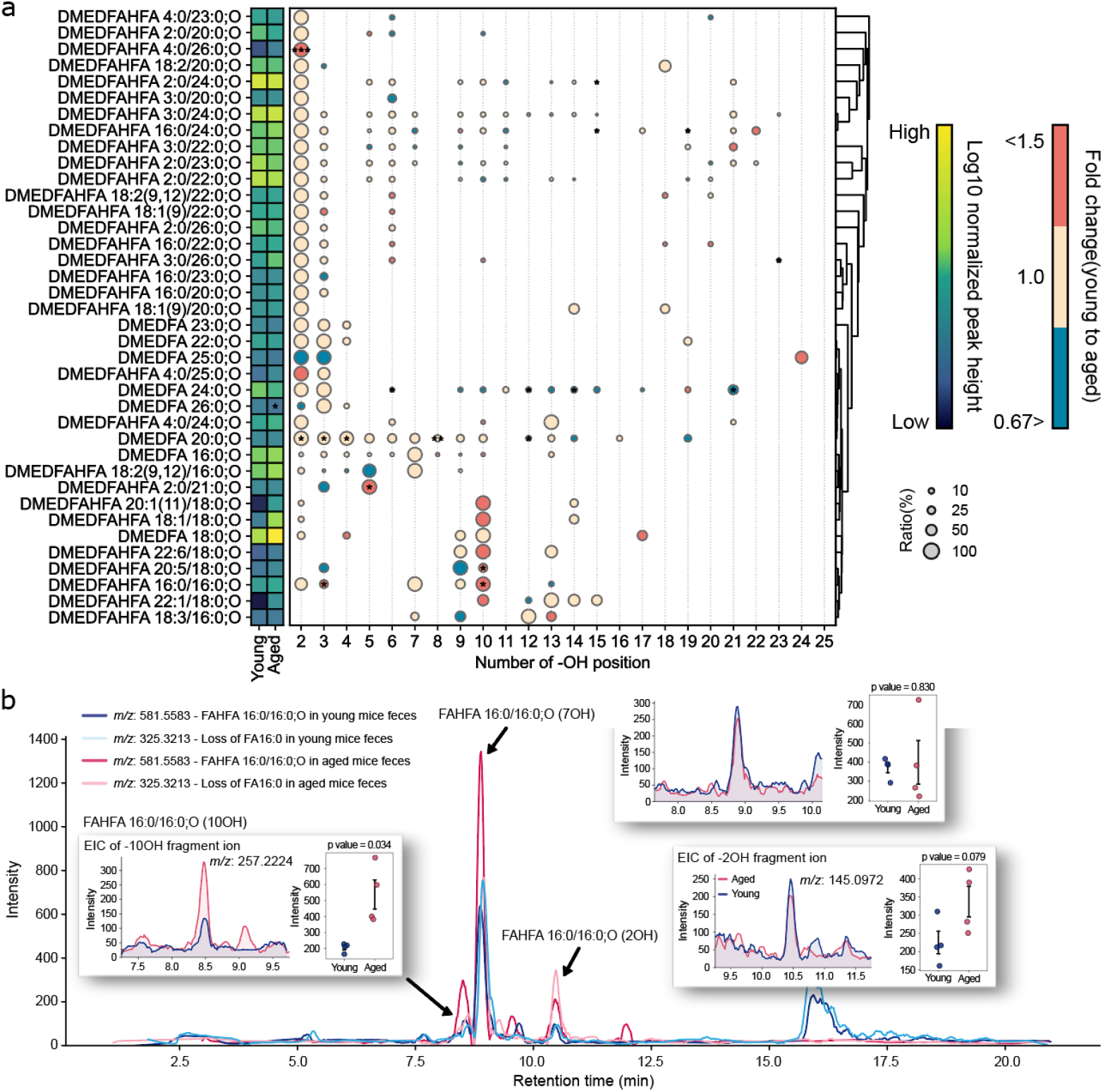
Profiling of fatty acid esters of hydroxy fatty acid (FAHFA) and HFA isomers based on the MS/MS chromatograms in SWATH-DIA-EAD-MS/MS data. (a) Distribution of OH positional isomers for 38 FAHFA and HFA species selected based on the signal-to-noise rations of >10 in precursor ion trace and the fact that the molecule contains saturated fatty acid as hydroxy fatty acid. Each row represents a representative lipid molecule, while columns indicate the OH position. The representative lipid name was characterized by EAD-DDA-MS/MS with MS-DIAL program. The color of each circle node reflects the fold changes in between aged (red) and young (blue) mice. Circle size represents the relative abundance of each OH position within a molecule (0–100%). The heatmap of the left panel shows the log2 value of normalized peak area in young and aged mice. A dendrogram on the right, constructed using Euclidean distance and complete linkage method based on the relative abundance values, illustrates the hierarchical clustering based on the lipid profile. The *p*-value was calculated by Welch’s t-test (two-sided, n=4). \**P* < 0.05, \*\**P* < 0.01, and \*\*\**P* < 0.001 (b) Showcase of MS/MS chromatograms for profiling FAHFA 16:0/16:0;O isomers. The product ion traces for precursor (*m/z* 581.5583) and diagnostic fragment of DMED-FAHFA (*m/z* 325.3213) in young and aged mice. The product ion traces for profiling 2OH (*m/z* 145.0972), 7OH (*m/z* 215.1754), and 10OH (*m/z* 257.2224) positional isomers of FAHFA 16:0/16:0;O; blue and red colors denote young and aged mice, respectively. The dot plots with *p*-value (student t-test, n=4) were also described.

We previously reported that SCFA/LHFA FAHFAs contain a hydroxy moiety at the 2OH position, which can be characterized by a unique fragment ion in ESI(−)-CID-MS/MS. Our results show that SCFA/LHFA-FAHFAs with SCFA 2:0 (acetic acid) as the branched chain tended to decrease with aging, whereas those with SCFA 3:0 (propionic acid) or SCFA 4:0 (butyric acid or isobutyric acid) as the branched chain tended to increase. In addition, this study using SWATH-DIA-EAD-MS/MS revealed that various OH-positional isomers existed, as judged by a signal-to-noise ratio greater than 10, and not only 2OH-LHFA, but also 3OH-LHFA were abundant in fecal samples (**Figure 5a**). Interestingly, while 3OH-FA having anti-inflammatory effects and agonist activity for peroxisome proliferator-activated receptor gamma (PPARγ) is abundant in fecal samples, the 3OH-form in FAHFAs was not highly abundant, suggesting that an enzymatic recognition system enriches 2OH-FAHFA in the gut microbiome^22^. Although several unusual HFAs with an OH moiety at the acyl chain terminal position have been detected, a recent study reported that omega-2 fatty acids have been detected in mammalian samples and confirmed using an ozone-induced dissociation technique^23^. The omega-2 fatty acids are potential substrates for terminal OH LHFA metabolites. In addition, our results showed that FAHFA 4:0/2O(FA 26:0) and FAHFA 16:0/10O(FA 16:0) significantly increased with age, whereas several FAHFAs such as FAHFA 2:0/3O(FA 21:0) and FAHFA 18:2/5O(FA 16:0) decreased with age (**Figure 5a and Figure 5b**).

## CONCLUSION

This paper describes a novel FAHFA profiling method using SPE and LC-SWATH-DIA-EAD-MS/MS with MS-DIAL updates. This screening method can be facilitated by considering the unique fragment ions at the OH position and acyl chain compositions. Moreover, this methodology is applicable for not only FAHFA, but also other fatty acid molecules such as HFAs. The MRMPROBS program for SWATH-DIA data allows the calculation of the ratio of OH-positional isomers that cannot be separated by conventional LC methods. Using the FAHFA analysis platform developed in this study, we characterized 85 FAHFA structural isomers in mouse feces. Our method is not limited to fecal samples, but can be applied to various organs and plasma, offering an environment to discover phenotype-associated FAHFA isomers. The gut microbiota produces a larger diversity of fatty acid metabolites than the host, many of which are still unknown. Although PNPLT2 is the response gene for FAHFA biosynthesis in host cells, the diversity and physiological activity of FAHFAs produced by gut bacteria remain largely unknown. Our method provides a deep understanding of the biological impact of FAHFAs and the complex metabolic network constructed by the host and microbiome.

## ASSOCIATED CONTENT

### Supporting Information

The Supporting Information is available free of charge on the ACS Publications website. Figure S1. Summary of fecal lipidome profiles before (left) and after (right) solid-phase extraction. Figure S2. Annotation rates of fatty acid esters of hydroxy fatty acid (FAHFAs) detected in feces lipid extracts from various enrichment conditions. Figure S3. Fragmentation pattern of DMED-derivatized hydroxy fatty acid (HFA). Table S1. Authentic standards of FAHFAs used in this study. Table S2. Lipidome table containing free fatty acids, hydroxy fatty acids, and FAHFAs. All raw LC-MS data and the related lipidome tables are available on the RIKEN DROP Met website (http://prime.psc.riken.jp) under index number DM0067. The Source Data for the figures was submitted. The MS-DIAL source code is available at https://github.com/systemsom-icslab/MsdialWorkbench.

## AUTHOR INFORMATION

## Author Contributions

Hiroshi T. designed this study. Y.K. performed LC-MS/MS analyses. Y.M., M.T., and H.T. developed MS-DIAL. Hiroaki T. improved the experimental design. M.H. and J.M. performed the mouse experiments. M.A. optimized the LC conditions. Y. K. and Hiroshi T. prepared the manuscript. All authors have thoroughly discussed this project and helped improve the manuscript.

## ACKNOWLEDGMENT

This study represents a portion of the dissertation submitted by Yuto Kurizaki to the Tokyo University of Agriculture and Technology in partial fulfillment of the requirements for his Ph.D. We thank Ms. Mie Honda for assistance with the LC-MS analysis. This study was supported by the JSPS KAKENHI (21K18216, H.T.), National Cancer Center Research and Development Fund (2020-A-9, H.T.), AMED Japan Program for Infectious Diseases Research and Infrastructure (21wm0325036h0001, H.T.), AMED Brain/MINDS (JP15dm0207001, H.T. and H.T.), JST National Bioscience Database Center (NBDC, H.T.), JST ERATO “Arita Lipidome Atlas Project” (JPMJER2101, M.A. and H.T.) and Technologically Advanced research through Marriage of Agriculture and engineering as Groundbreaking Organization (TAMAGO to J.M. and Hiroshi T.).

## Notes

### Competing Interest Statement

The authors have declared no competing interest.

## REFERENCES

1. Brejchova, K., et al. Understanding FAHFAs: From structure to metabolic regulation. Prog Lipid Res 79, 101053 (2020).

2. Yore, M.M., et al. Discovery of a class of endogenous mammalian lipids with anti-diabetic and anti-inflammatory effects. Cell 159, 318–332 (2014).

3. Patel, R., et al. ATGL is a biosynthetic enzyme for fatty acid esters of hydroxy fatty acids. Nature 606, 968–975 (2022).

4. Aryal, P., et al. Distinct biological activities of isomers from several families of branched fatty acid esters of hydroxy fatty acids (FAHFAs). J Lipid Res 62, 100108 (2021).

5. Riecan, M., Paluchova, V., Lopes, M., Brejchova, K. & Kuda, O. Branched and linear fatty acid esters of hydroxy fatty acids (FAHFA) relevant to human health. Pharmacol Therapeut 231, 107972 (2022).

6. Gowda, S.G.B., et al. Identification of short-chain fatty acid esters of hydroxy fatty acids (SFAHFAs) in a murine model by nontargeted analysis using ultra-high-performance liquid chromatography/linear ion trap quadrupole-Orbitrap mass spectrometry. Rapid Commun Mass Sp 34, e8831 (2020).

7. Yasuda, S., et al. Elucidation of Gut Microbiota-Associated Lipids Using LC-MS/MS and 16S rRNA Sequence Analyses. Iscience 23, 101841 (2020).

8. Tsugawa, H., et al. A lipidome atlas in MS-DIAL 4. Nat Biotechnol 38, 1159–1163 (2020).

9. Kuda, O., et al. Docosahexaenoic Acid-Derived Fatty Acid Esters of Hydroxy Fatty Acids (FAHFAs) With Anti-inflammatory Properties (vol 65, pg 2580, 2016). Diabetes 65, 3516–3516 (2016).

10. Randolph, C.E., Marshall, D.L., Blanksby, S.J. & McLuckey, S.A. Charge-switch derivatization of fatty acid esters of hydroxy fatty acids via gas-phase ion/ion reactions. Anal Chim Acta 1129, 31–39 (2020).

11. Ding, J., et al. In-Silico-Generated Library for Sensitive Detection of 2-Dimethylaminoethylamine Derivatized FAHFA Lipids Using High-Resolution Tandem Mass Spectrometry. Anal Chem 92, 5960–5968 (2020).

12. Zhu, Q.F., Yan, J.W., Zhang, T.Y., Xiao, H.M. & Feng, Y.Q. Comprehensive Screening and Identification of Fatty Acid Esters of Hydroxy Fatty Acids in Plant Tissues by Chemical Isotope Labeling Assisted Liquid Chromatography-Mass Spectrometry. Anal Chem 90, 10056–10063 (2018).

13. Takeda, H., Takeuchi, M., Hasegawa, M., Miyamoto, J. & Tsugawa, H. A Procedure for Solid-Phase Extractions Using Metal-Oxide-Coated Silica Column in Lipidomics. Anal Chem 96, 17065-17070 (2024).

14. Tsugawa, H., et al. MRMPROBS: A Data Assessment and Metabolite Identification Tool for Large-Scale Multiple Reaction Monitoring Based Widely Targeted Metabolomics. Anal Chem 85, 5191–5199 (2013).

15. Tsugawa, H., Kanazawa, M., Ogiwara, A. & Arita, M. MRMPROBS suite for metabolomics using large-scale MRM assays. Bioinformatics 30, 2379–2380 (2014).

16. Takeda, H., et al. MS-DIAL 5 multimodal mass spectrometry data mining unveils lipidome complexities. Nat Commun, 10.1038/s41467-41024-54137-w (2024).

17. Tsugawa, H., et al. Mass Spectrometry Data Repository Enhances Novel Metabolite Discoveries with Advances in Computational Metabolomics. Metabolites 9, 119 (2019).

18. Kiuchi, S., et al. Using Variable Data-Independent Acquisition for Capillary Electrophoresis-Based Untargeted Metabolomics. J Am Soc Mass Spectr 35, 2118–2127 (2024).

19. Tokiyoshi, K., et al. Using Data-Dependent and - Independent Hybrid Acquisitions for Fast Liquid Chromatography-Based Untargeted Lipidomics. Anal Chem 96, 991–996 (2024).

20. Tsugawa, H., et al. MS-DIAL: data-independent MS/MS deconvolution for comprehensive metabolome analysis. Nat Methods 12, 523–526 (2015).

21. Folz, J., et al. Human metabolome variation along the upper intestinal tract. Nat Metab 5, 777–788 (2023).

22. Pujo, J., et al. Bacteria-derived long chain fatty acid exhibits anti-inflammatory properties in colitis. Gut 70, 1088–1097 (2021).

23. Menzel, J.P., et al. Ozone-enabled fatty acid discovery reveals unexpected diversity in the human lipidome. Nat Commun 14, 3940 (2023).

