## Supporting information for "Data-independent acquisition coupled with electron-activated dissociation for in-depth structure elucidation of fatty acid ester of hydroxy fatty acids"


### Contents


Supplementary Figures 1, 2, and 3

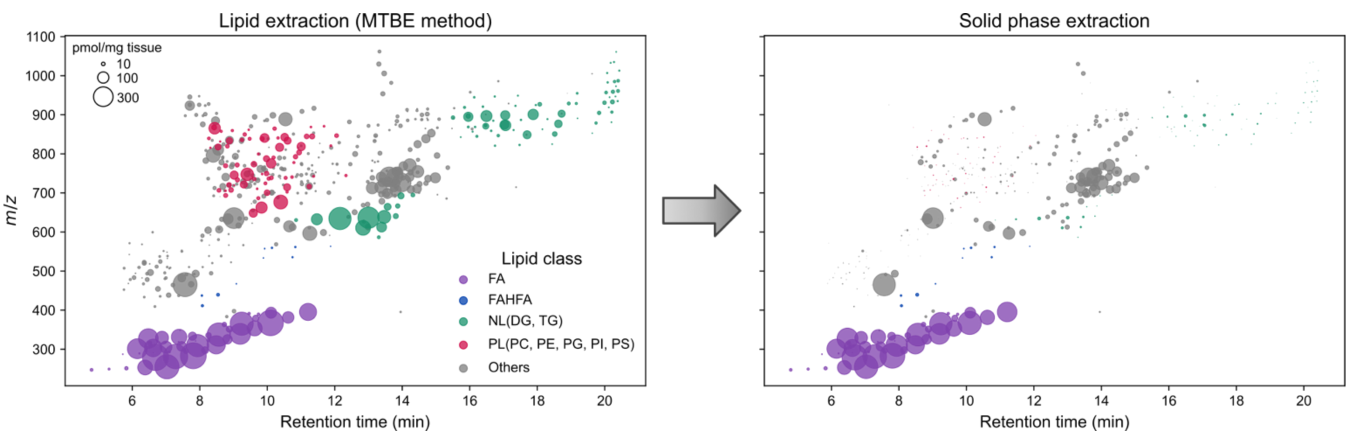


**Supplementary Figure 1. Summary of fecal lipidome profiles before (left) and after (right) solid-phase extraction.** The x- and y-axis denote retention time (min) and *m/z*, respectively. Point size indicates the normalized ion abundance normalized by internal standards where the unit can be interpreted as pmol/mg tissue. Colors represent lipid classes: FA (purple), FAHFA (blue), neutral lipid (NL; DG, TG) (green), phospholipid (PL; PC, PE, PG, PI, PS) (red), and the others (gray).

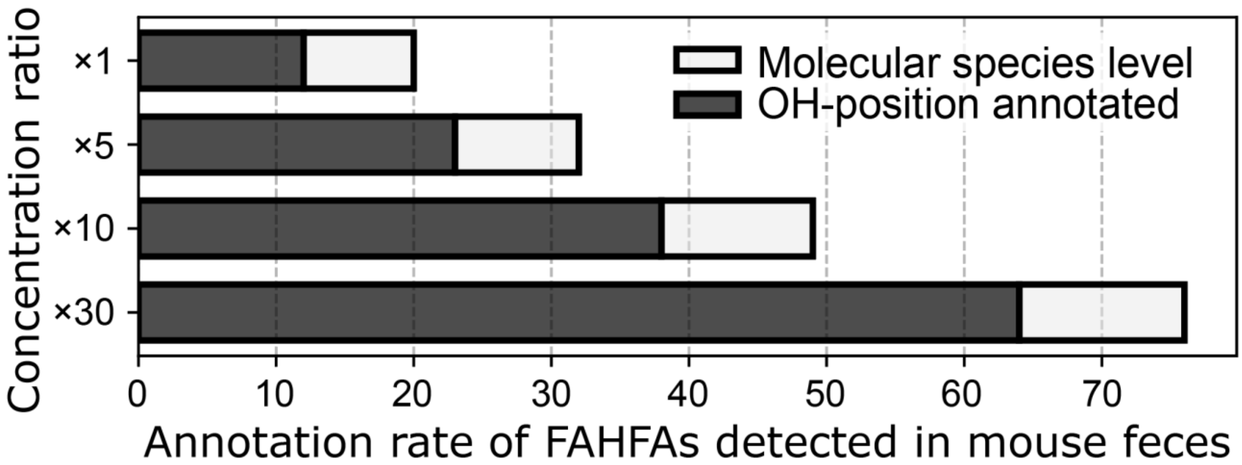


**Supplementary Figure 2. Annotation rates of fatty acid esters of hydroxy fatty acid (FAHFAs) detected in feces lipid extracts from various enrichment conditions.** The x-axis represents the annotation rate of FAHFAs, whereas the y-axis represents the concentration factors (1×, 5×, 10×, and 30×) of the lipid extracts. The term of "x1" means the original concentration where the lipid extract from 6.7 mg of mouse feces is dissolved in 100 µL of methanol in LC-MS vial. is White and green bars indicate the numbers of annotated FAHFAs at the molecular species- and OH-resolved levels, respectively.

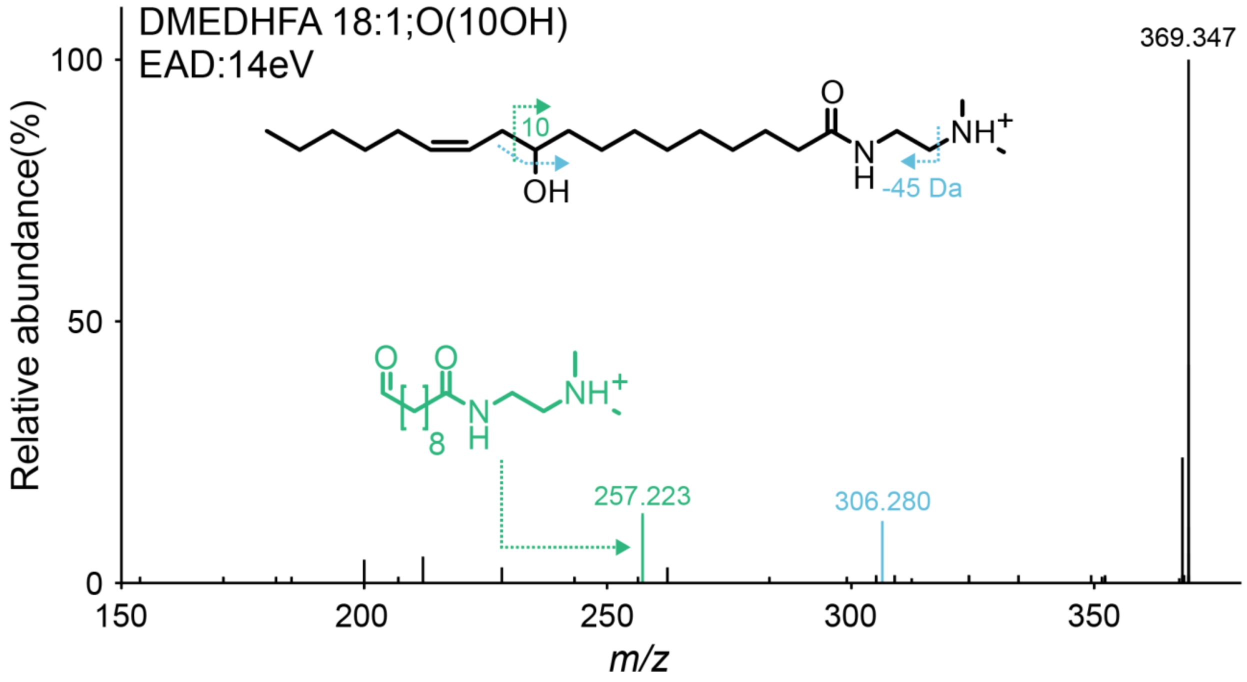


**Supplementary Figure 3. Fragmentation pattern of DMED-derivatized hydroxy fatty acid (HFA).** The product ion spectrum of DMED-derivatized 10-hydroxy-12(Z)-octadecenoic acid was described.
